# Designing a Novel Multi-Epitope Vaccine against SARS-CoV-2; Implication for Viral Binds and Fusion Inhibition through Inducing Neutralizing Antibodies

**DOI:** 10.1101/2021.06.16.448772

**Authors:** Seyed Amir Hossein Mohammadzadeh Hosseini Moghri, Mojtaba Ranjbar, Hadi Hassannia, Fatemeh Khakdan

## Abstract

Recently the prevalence of severe acute respiratory syndrome coronavirus 2 (SARS-CoV-2) has become a pervasive threat to public health so it is an emergency to vaccine development. The SARS-CoV-2 spike (S) glycoprotein plays a vital role in binds and fusion to the angiotensin-converting enzyme 2 (ACE2). The multi-epitope peptide vaccines are capable of inducing the specific humoral or cellular immune responses. In this regard, the RBD and spike cleavage site is the most probable target for vaccine development to inducing binds and fusion inhibitors neutralizing antibodies. In the present study, several immunoinformatics tools are used for analyzing the spike (S) glycoprotein sequence including the prediction of the potential linear B-cell epitopes, B-cell multi-epitope design, secondary and tertiary structures, physicochemical properties, solubility, antigenicity, and allergenicity for the promising vaccine candidate against SARS-CoV-2.

## 1. Introduction

SARS-CoV-2, as a virus firstly discovered in Wuhan, China, in December 2019, is a new coronavirus in humans (1, 2, 3) that can cause severe acute respiratory syndromes. The nuclear material of the SARS-CoV-2 is positive-sense single-stranded RNA (+ssRNA) that possess viral RNA genomes of about 30 Kbp (4). The SARS-CoV-2 genomes consist of a 5′ cap structure together with a 3′ poly (A) tail, which allows it to acts as mRNA for translation of the replicase polyproteins (5). The genome of SARS-CoV-2 encodes four structural proteins that could induce a much higher T cell response than the non-structural proteins (nsps) (6). The structural proteins possess various roles in viral processes, such as the formation of the virus particles including spike (S) glycoprotein, envelope (E) glycoprotein, membrane (M) glycoprotein, and nucleoprotein (N) (7, 8). SARS-CoV-2 initiates the infection of the host cells mediated by the spike (S) glycoprotein, which is found in the homotrimers protruding from the viral surface (9). The S glycoprotein is cleaved by the host cells’ furin-like protease and TMPRSS2 (Type II transmembrane serine proteases) (10, 11) into two functional subunits including S1 and S2. Subunit S1 contains the RBD that is a binding agent for the peptidase domain (PD) of ACE2 as the host cell receptor. Meanwhile, S2 functions are the viral-cellular membranes fusion (8, 9). The ACE2 is a type-I membrane receptor that acts as a peptide hormone. This enzyme plays the primary physiological role in the maturation of angiotensin (Ang) and is responsible for vasoconstriction and blood pressure. ACE2 is expressed in the heart, lungs, kidneys, and intestine (9). As the RBD of the S1 glycoprotein is surface-exposed and also as a critical protein that mediates entry into host cells, it could serve as the main target to induce neutralizing antibodies (nAbs) to diagnostic immune people and therapeutic vaccine application (8). The multi-epitope peptide-based vaccine, thanks to several advantages including supreme specificity, suitable safety, simplicity in production, storage, and stability, has become an attractive area in the field of vaccine research (12). Therefore, spikes can be considered as the main target to inducing the neutralizing antibodies as binding and indirectly fusion inhibitors for the vaccine design both immune people diagnostics and vaccine application.

## 2. Materials and Methods

### 2.1. RBD Sequence Retrieval and Structural Prediction

The amino acid sequences of the RBD with the accession number of QHD43416.1 were retrieved from the NCBI database in FASTA format (13). The spike (S) as a transmembrane glycoprotein has a molecular weight of about 150 kDa (7).

### 2.2. Linear B-cell epitope prediction

To initiate an immune response, first, the B cell epitope interacts with B lymphocytes (14). ABCpred Prediction Server (http://www.imtech.res.in/raghava/abcpred/) was applied to predict potential linear Based on the recurrent neural networks (RNNs), B-cell epitopes were predicted for 14-mers with a default threshold of 0.51 (15).

### 2.3. B-cell multi-epitope vaccine candidate design

Vaccine candidate constructed based on six surface-exposed linear B-cell epitopes in the way that epitopes fused via AAY linkers (16). Furthermore, after the RBD binding with ACE2, spike was cleaved into S1 and S2 subunit in the 680-SPRRAR-685 residue by host proteases at S1/S2 cleavage sites such as furin and TMPRSS2 and new conformation can facilitate viral fusion with the host cell membrane (10, 11). Thus can induces the neutralizing antibodies as indirect fusion inhibitors due to prohibit of conformation change.

### 2.4. Physiochemical properties, solubility, antigenicity and allergenicity prediction

Various physicochemical features of the RBD, including theoretical pI, instability index, aliphatic index, estimated half-life in the mammalian reticulocytes *in vitro*, extinction coefficient, grand average of hydropathicity (GRAVY), and molecular weight, were determined using the online web server ProtParam (http://web.expasy.org/protparam/) (17). Solubility in the water of the RBD was evaluated using the protein-sol server (https://protein-sol.manchester.ac.uk/) (18). The peptide antigenicity characteristics were analyzed via VaxiJen v2.0 as an online antigen prediction server (http://www.ddgpharmfac.net/vaxijen/). The threshold for virus models was set at ≥ 0.4 (19). The web-based servers AllergenFP v.1.0 (http://www.ddg-pharmfac.net/AllergenFP/) (20) were used for predicting the epitope allergenicity.

### 2.5. Secondary structure prediction

The secondary structure of the final peptide with position-specific iterated BLAST (PSI-blast) was predicted based on secondary structure PREDiction (PSIPRED) (http://bioinf.cs.ucl.ac.uk/psipred/) (21). Also, the secondary structure properties of the RBD were predicted via the RaptorX Property web server (http://raptorx.uchicago.edu/StructurePropertyPred/predict/) (22).

### 2.6. Tertiary structure prediction and validation

The three-dimensional (3D) structure of the spike (S) glycoprotein was downloaded from the Iterative Threading ASSEmbly Refinement (I-TASSER) server (https://zhanglab.ccmb.med.umich.edu/I-TASSER/). Also, we generated RBD using I-TASSER. Through multiple threading alignments and iterative structural assembly simulations, I-TASSER first generates three-dimensional (3D) atomic models starting from an amino acid sequence (23). Next, visualization and analyses were performed by building a 3D model in a PDB format using PyMOL (24). The built model was validated as a vital step to detect potential errors in predicted 3D models (16). Subsequently, a Ramachandran plot was generated through the Ramachandran Plot server (https://zlab.umassmed.edu/bu/rama/) to represent energetic visualization of allowed and disallowed dihedral angles psi (ψ) and phi (ϕ) of amino acid. These angles were then calculated based on the van der Waal radius of the side chain. The Ramachandran Plot results demonstrate the percentage of residues in allowed and disallowed regions that define the quality of the modeled structure (25).

### 2.7. Codon adaptation and *in silico* cloning of the designed vaccine

The JCat server (Java Codon adaptation tool) (http://www.prodoric.de/JCat/) was performed for codon optimization, to the multi-epitope construct expression in pET-28a(+) vector (26). Codon optimization leads to the higher expression rate of the final vaccine in the E. coli K12 as an expression host because the codon usage of human and selected host differs from each other. Three additional options were applied to avoid the rho-independent transcription termination, prokaryote ribosome binding site, and cleavage site of restriction enzymes for increasing the efficiency of a translation process. The JCat result includes the codon adaptation index (CAI) and percentage GC content, which can be used to evaluate protein expression levels. CAI provides information on codon usage biases; the ideal CAI score is 1.0 but greater than 0.8 is considered a good score (27). The GC content of a sequence should range between 30–70%, outside this range of GC content values considered unfavorable effects on translational and transcriptional efficiencies (28). To clone the optimized final vaccine sequence the HindIII and SalI restriction sites were added at the N- and C- terminal sites of the final construct, respectively. Finally, the optimized sequence (with added restriction sites) was inserted into the pET-28a(+) vector using the SnapGene restriction and insertion cloning module to ensure vaccine expression.

## 3. Results

### 3.1. RBD Sequence analysis and Structural Prediction

The amino acid residues of the RBD including 254 amino acids with residues ranging from 330 to 583 were selected. I-TASSER generated 3D-model was used to represent the RBD residues of trimeric spike (S) glycoprotein (Figure 1A) based on the surface exposed and S-ACE2 (Figure 1B) using PyMOL.

**Figure 1.**
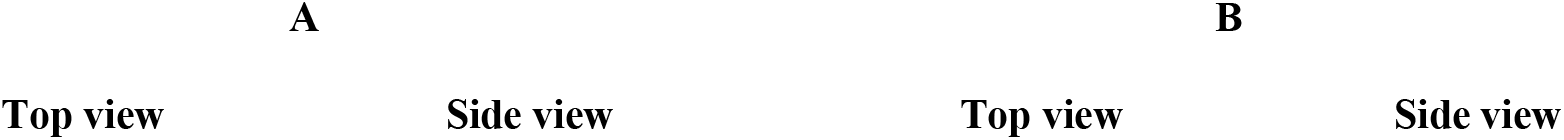
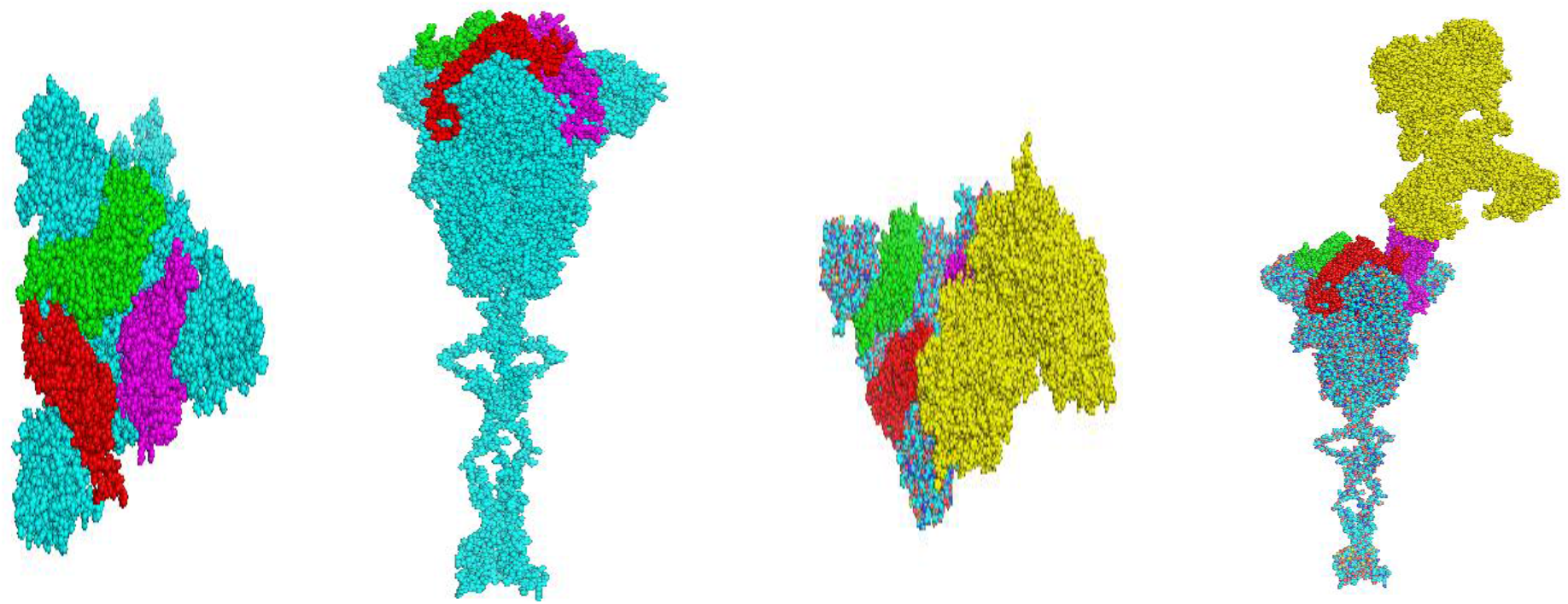
3D-model of trimeric spike (S) glycoprotein and it’s binding to ACE2 by using PyMOL: **a**) 3D-model of the RBD of chains A (red), B (green), C (magenta) and spike (S) glycoprotein residues shown in cyan and **b**) Interaction of S-ACE2 (yellow)

### 3.2. Linear B-cell epitope prediction

The ABCpred server predicts 25 Linear B cell epitopes, of which 18 had a significant VaxiJen score. However, among the 18 predicted linear B cell epitopes, 7 epitopes, including “APGQTGKIADYNYK”, “KCYGVSPTKLNDLC”,“VKNKCVNFNFNGLT”,“GVGYQPYRVVVLSF”,“NFNFNGLTGTGVLT”, “PYRVVVLSFELLHA”, and “PTKLNDLCFTNVYA” have the VaxiJen score of more than 1. This suggests the high antigenicity nature of these epitopes and could be considered as the most potential antigenic B cell epitopes that would interact with B lymphocytes and induce neutralizing antibodies (Table 1 and Figure 2).

**Table 1.**
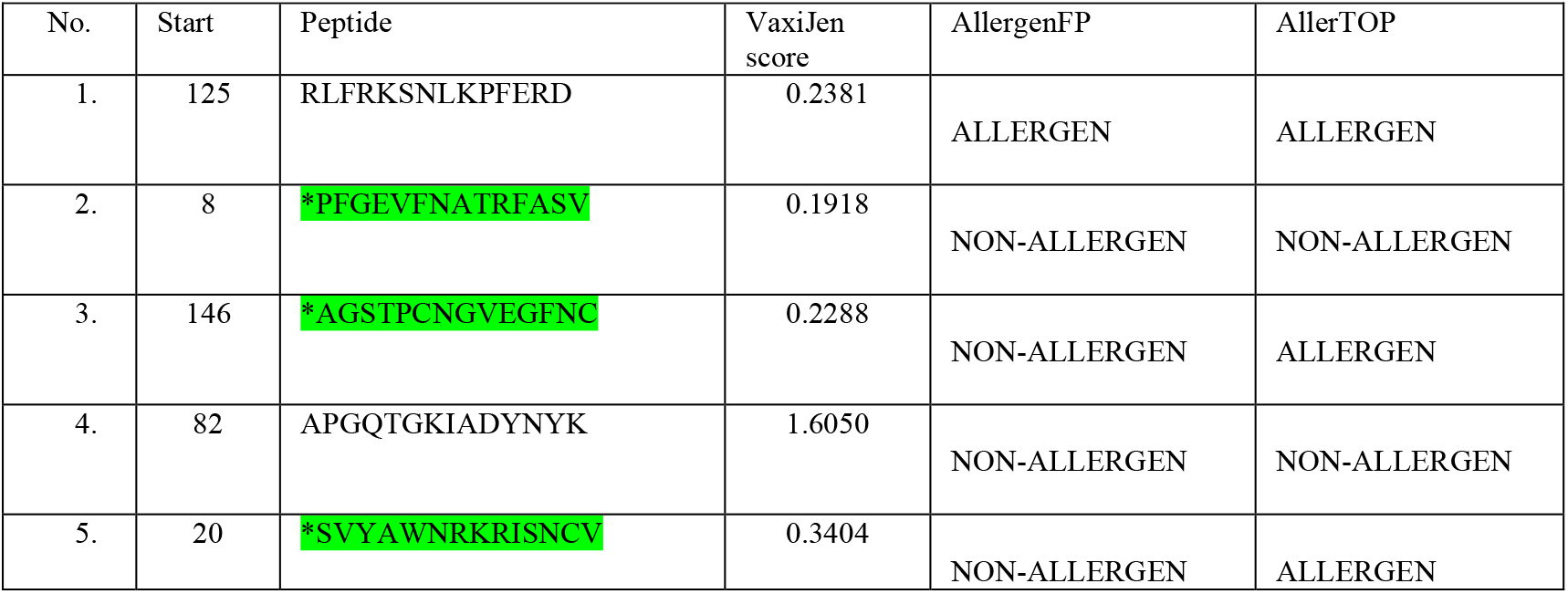
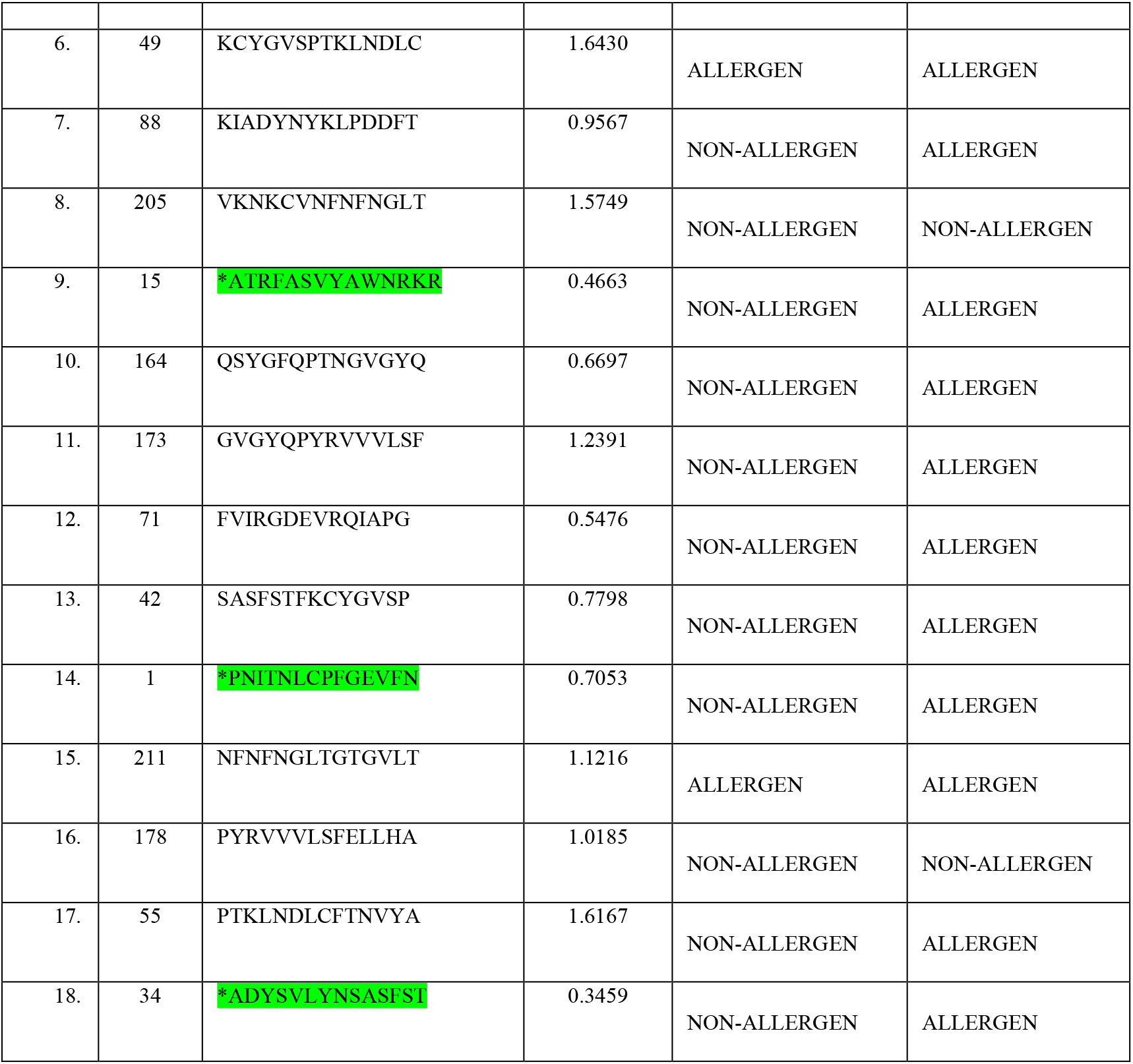
Predicted linear B cell epitopes

**Figure 2.**
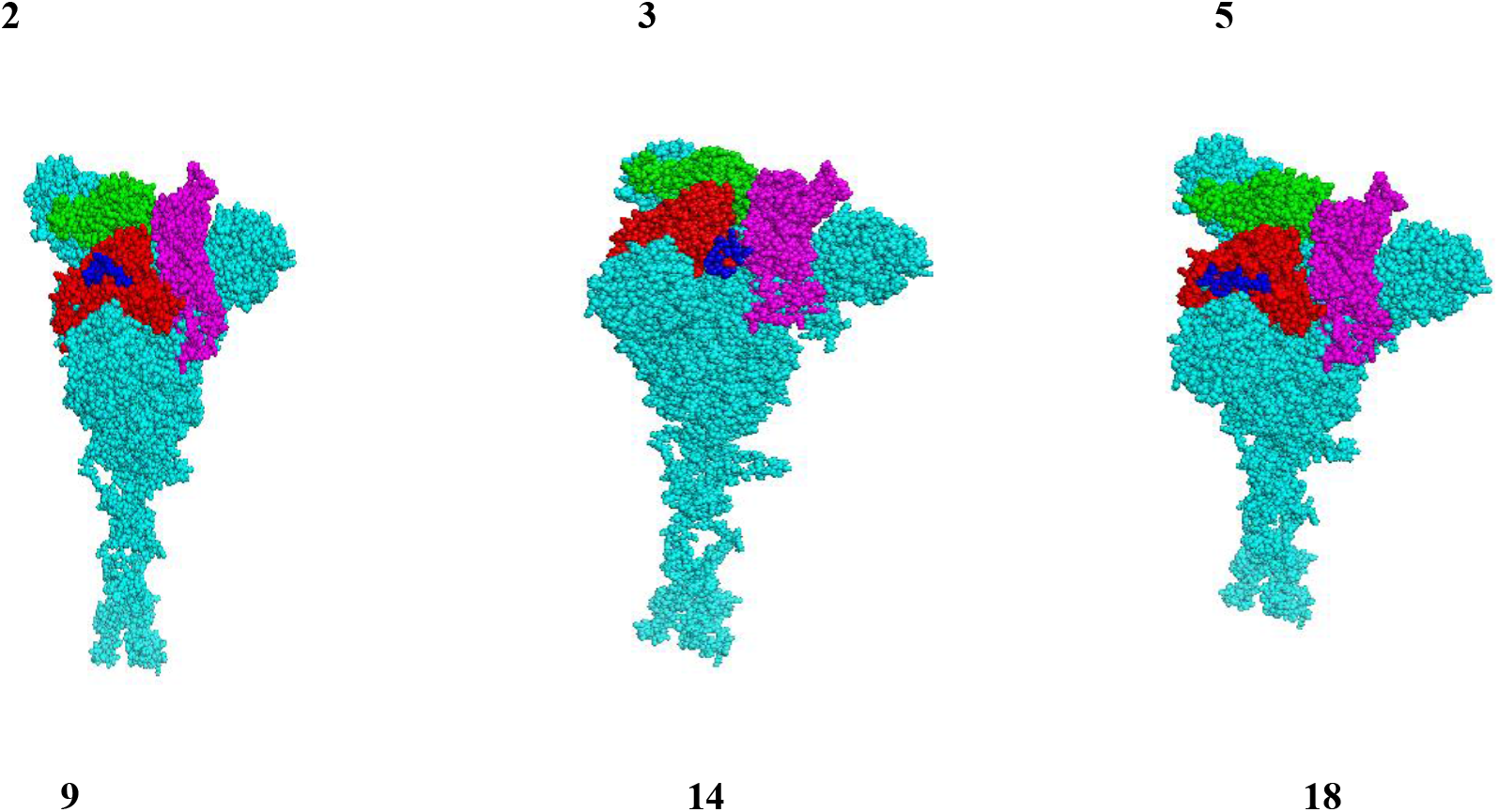
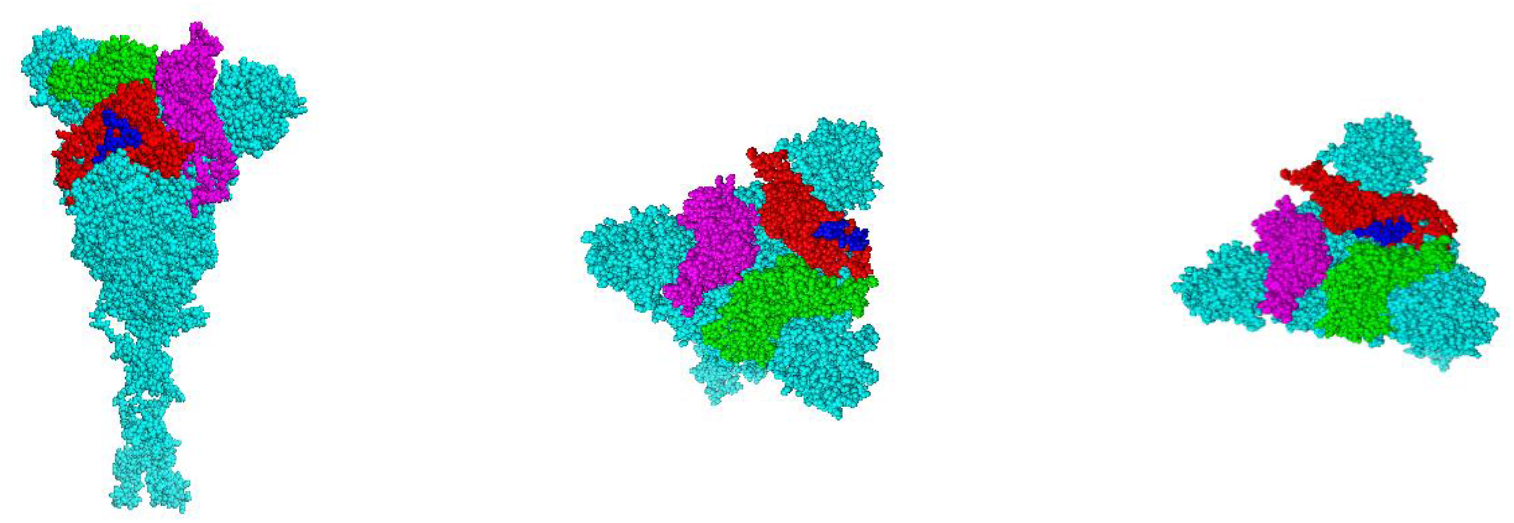
3D-model of the receptor-binding domain (RBD) of chain A (red), B (green), C (magenta) of spike (S) glycoprotein residues shown in cyan and the views of six surface-exposed linear B-cell epitopes of the RBD presented in blue on one of the chains (figures numbers is according to table’s numbers).

### 3.3. B-cell multi-epitope Vaccine design

To generating the B-cell multi epitopes construct, we fused six epitopes with AAY linkers. Furthermore, the 680-SPRRAR-685 residue targeting by host proteases at S1/S2 cleavage site was added at the N-terminal by AAY linkers too. Finally to protein purification, a 6x His tag was added at the C-terminal of the final vaccine and final sequence fixed between the start and stop codon. The final vaccine constructed with 114 residues long derived from 7 merged peptide sequences, a schematic diagram of the vaccine is presented in figure 3.

**Figure 3.**
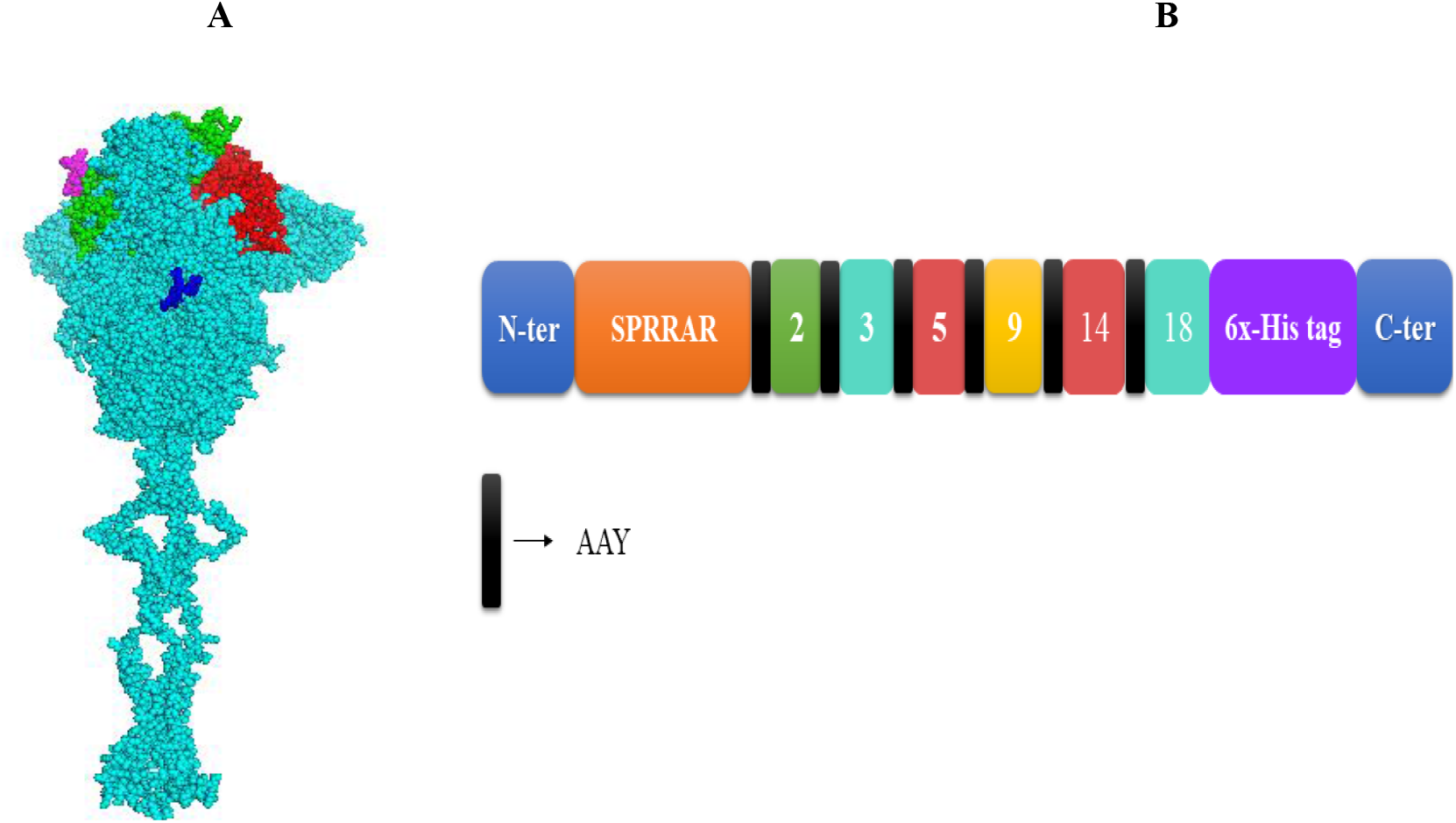
The vaccine designed has a 114 in length (A) presented the target residue (680-SPRRAR-685) of host proteases at the S1/S2 cleavage site in the chain A of spike (S) glycoprotein. (B) Schematic presentation of the final multi-epitope vaccine construct containing a protease targeting site (orange) in N-terminal linked with 6 epitopes while fused together through an AAY linker (black).

### 3.4. Physiochemical properties, solubility, antigenicity and allergenicity prediction

The vaccine candidate had an estimated molecular weight of about 12.6 kDa, the theoretical isoelectric point (pI) of 9.51 (due to the vaccine’s alkaline nature), instability index of 39.63 (the stability of the candidate vaccine), the grand average of hydropathicity (GRAVY) index of −0.268 (i.e., hydrophilic protein). Also, the aliphatic index was 53.33, suggesting that the vaccine may be stable for a wide temperature range. Moreover, estimated half-life in the yeast, escherichia coli in vivo and mammalian reticulocytes in vitro was >20, >10 and, 1.9 hours respectively. Vaccine solubility in water was 0.61 and its extinction coefficient was calculated to be from 25900 to 26150 M^−1^cm^−1^. So, it can provide support in the quantitative study of protein-ligands and protein-protein interaction in solution. The results of physicochemical properties and also the antigenicity and allergenicity of candidate peptides are shown in Table 2.

**Table 2.**
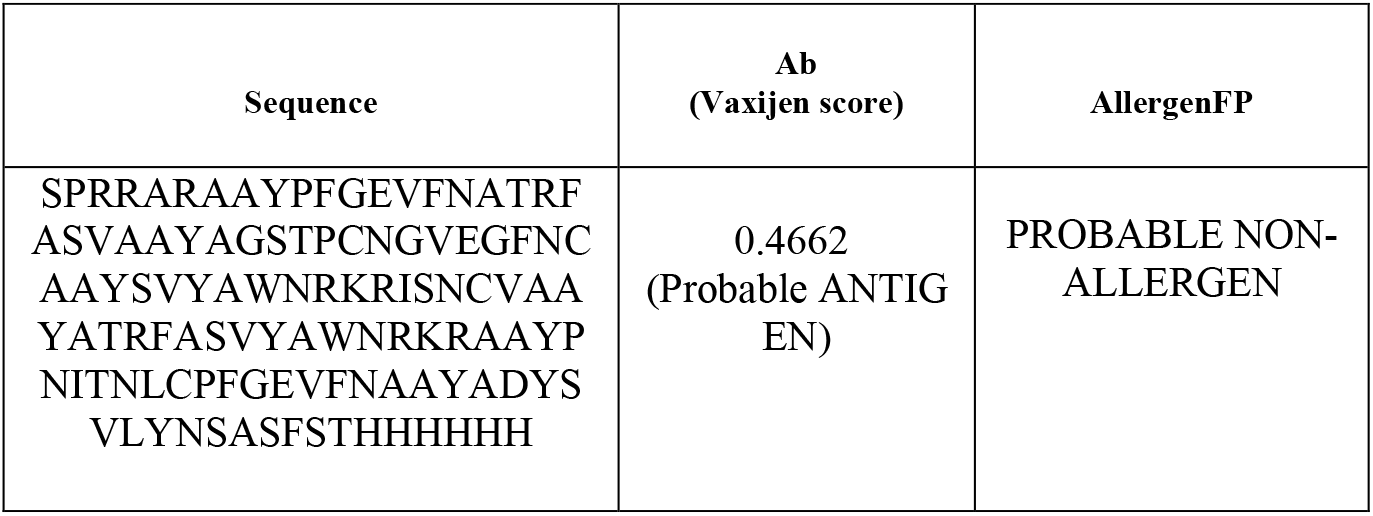
The antigenicity and the allergenicity of the vaccine.

### 3.5. Secondary structure prediction

The predicted PSIPRED secondary structure of the final chimeric peptide, which was obtained from PSI-BLAST, was submitted in FASTA format. Overall, the predicted secondary structure revealed that the protein has 24% alpha-helix, 7% beta-strand, and 68% coil (Fig. 4A). Besides, taking into account the amino acids solvent accessibility, it was predicted that 43, 21, and 34% of the acids were exposed, medium exposed, and buried, respectively. The RaptorX Property server predicted a total of 17 residues (14%) to be located in disordered domains.

**Figure 4.**
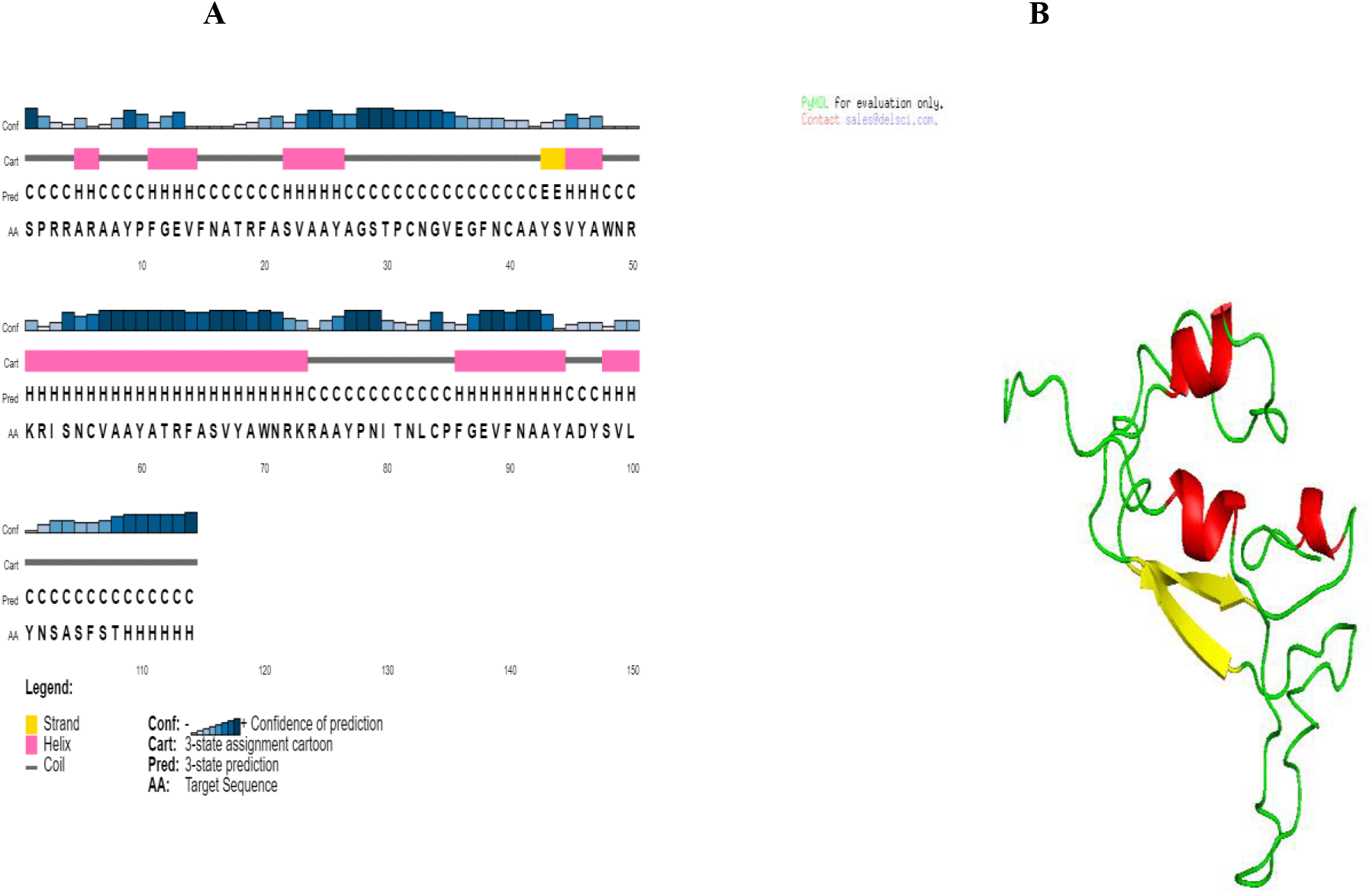
Structure prediction of the final peptide: **a**) PSIPRED predicted secondary structure, b) predicted 3D model homology structure of the RBD (helix, sheet, and loop are represented by red, yellow, and green, respectively)

### 3.6. Tertiary structure model and validation

Five tertiary structure models of the final peptide were predicted based on 10 threading templates by the I-TASSER server. The Z-score values of the top 10 chosen templates are ranging from 1.48 to 2.33, indicating a good alignment. The C-score values of five predicted models range from −2.26 to −3.84. The optimal range of the C-score value is between −5 and 2. The model with the highest C-score was selected via homology modeling (Fig. 4B). This model had an estimated TM-score and RMSD of 0.45±0.14 and 9.1±4.6Å, respectively. The TM-score has been suggested as a scale for measuring the structural similarity between two structures. Here, a TM-score of more than 0.5 indicates a correct topology model (28). Ramachandran Plot tool was used for evaluating the reliability model, plot generation Ramachandran, and determine the energy of the stable conformation of the psi (ψ) and phi (Φ) twisting or dihedral angles for every amino acid. The results of the Ramachandran plot analysis shows that the number of residues in the favored region was 81.37%. Also, the number of residues in the allowed region was 16.66%, and only 1.96% of the residues were found in the outlier region (Figure 5).

**Figure 5.**
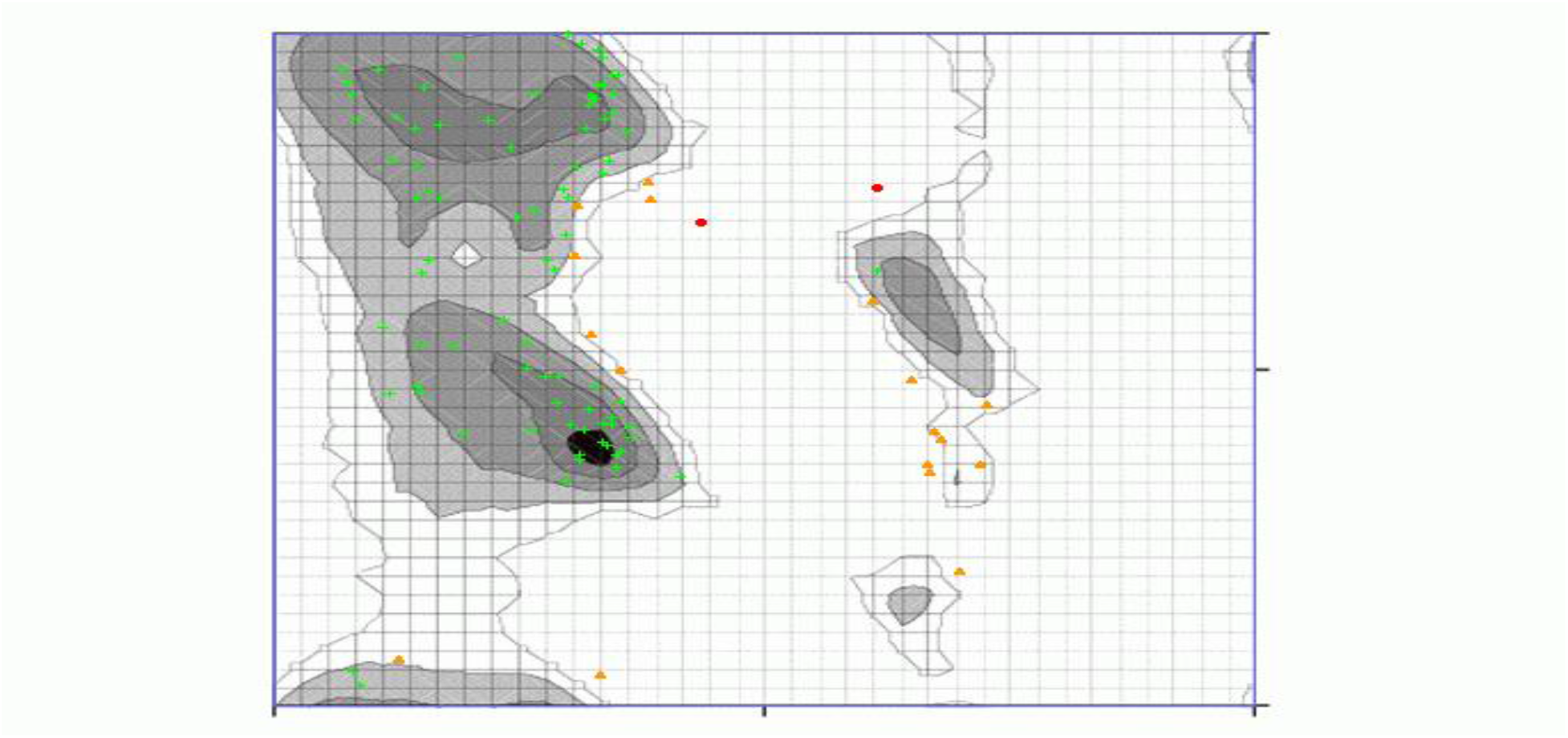
Tertiary structure modeling and validation by Ramachandran plot of the final vaccine modeled

### 3.6. Codon adaptation and *in silico* cloning of the designed vaccine

The Java Codon adaptation tool (JCat) was utilized to optimize codon usage of the constructed vaccine to Escherichia coli (strain K12) for maximal protein expression and different the expression system of the human and E. coli. The optimized codon sequence has 354 nucleotides in length. The Codon Adaptation Index (CAI) of the optimized codon sequence was discovered to be 0.96. The optimal range of CAI index is between zero to one, showing the likely success of the target gene expression and the sequence GC content was 52.3% which is also satisfactory due to the GC content optimal percentage range is between 30% and 70% showing the possibility of good expression in the E. coli host. Eventually, for restriction cloning of the final vaccine, designed sequence was constructed by adding into the pET28a(+) vector between HindIII and SalI restriction sites at the N-and C- terminal sites, respectively (Figure 6).

**Figure 6.**
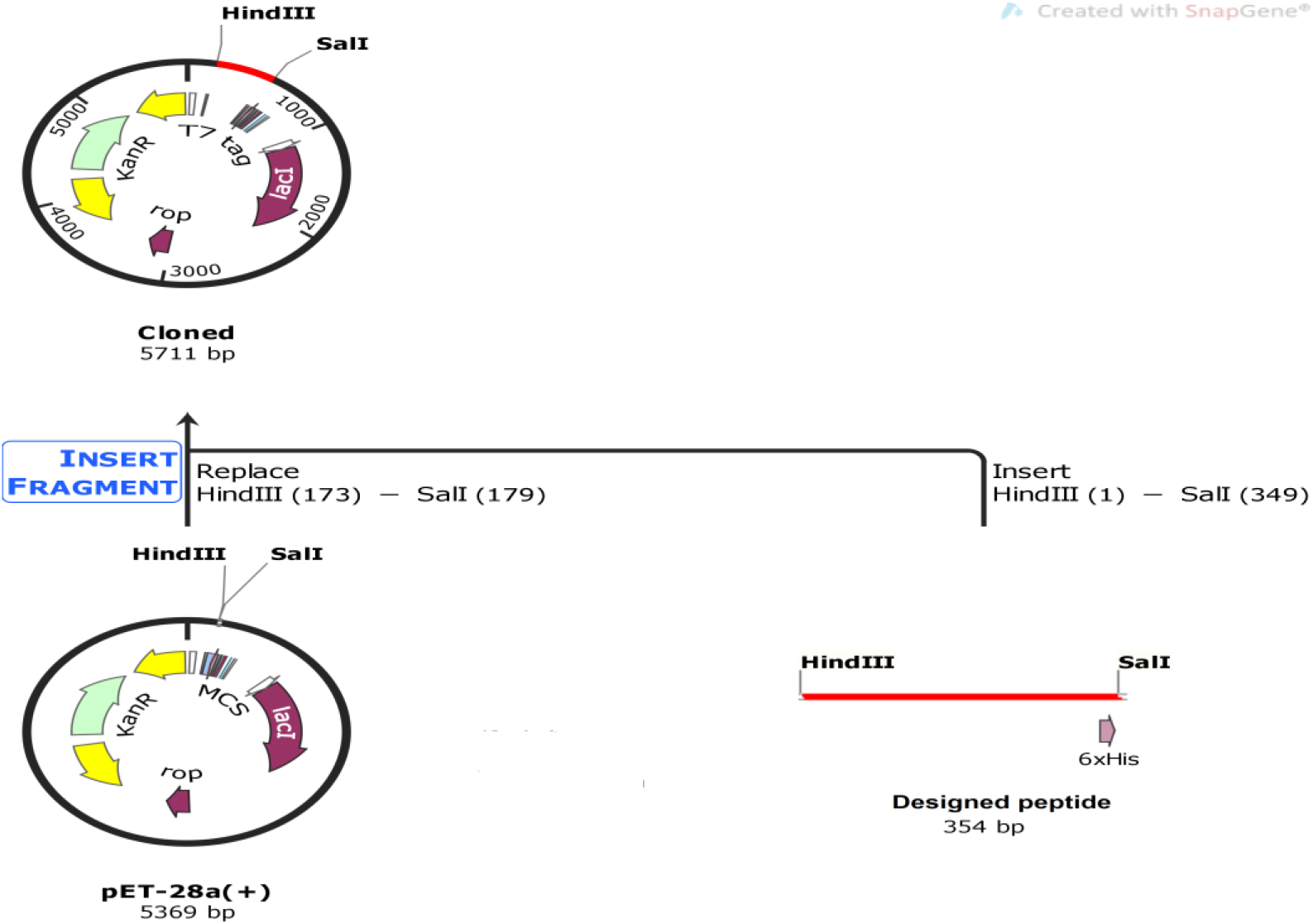
The final vaccine *in silico* restriction and insertion cloning into the pET28a(+) expression vector where the magenta part indicates the construct gene coding and the vector backbone represented in the black circle. The His-tag is located at the carboxy-terminal end.

## 4. Discussion

Recently the prevalence of severe acute respiratory syndrome coronavirus 2 (SARS-CoV-2) has become a pervasive threat to public health so it is an emergency to vaccine development (1). As mentioned the RBD protein located in the S1 subunit is responsible for binds to the ACE2 receptor and then due to S1/S2 cleavage by host cell proteases the conformation changed which led to fusion the virus membrane with the host cell membrane can be led to the entry of the virus into the host cells. Accordingly, spikes serve as the main target to induce the neutralizing antibodies as binds and fusion inhibitors for the vaccine design both immune people diagnostic and therapeutic application against SARS-CoV-2. Recently the multi-epitope vaccine designing is an emerging field that fortunately has not only demonstrated in vivo efficacy with protective immunity (29, 30, 31) but also arrived phase-I clinical trials (32, 33, 34, 35). In like manner, the immunoinformatic procedure of vaccine designing has recently been used for designing multi-epitope vaccines against Nipah virus (36), Dengue (28), Malaria (37) and Hendra virus (38). In this current study, we constructed six surface-exposed linear B-cell epitopes with AAY linkers. Furthermore, the 680-SPRRAR-685 residue targeting by host proteases at S1/S2 cleavage site was added at the N-terminal by AAY linkers too and we applied various immunoinformatics tools to gather information about the candidate vaccine. The vaccine has been shown vaxijen scores 0.46 and also the absence of allergenic properties could be promising potential as a vaccine candidate. As a result, the vaccine molecular weight is about 12.6 kDa and the predicted vaccine solubility was 0.61 indicates good solubility due to any scaled more than 0.45 is predicted to have a higher solubility (18). Moreover, the predicted instability index shows the stability upon expression of the candidate vaccine and due to the GRAVY it can interact in aqueous solutions (28). Overall the results of physicochemical properties and also the antigenicity and allergenicity of candidate peptide are satisfactory. Considering the importance of secondary and tertiary structures knowledge in target protein as a key factor in vaccine design (39), and results demonstrated that the vaccine contained mainly of coils (68%), with only 14% of residues disordered. The 3D structure of the vaccine demonstrated favorable features based on the Ramachandran plot which indicates that the total percentage of favored and allowed region residues is 98.03%. As exceeding 90% is an ideal result, the proposed model can be considered reliable which shows satisfaction about the quality of the overall model (25). The codon optimization was performed to attain the high-level expression of our designed vaccine in E. coli (strain K12). Both the CAI (0.96) and the GC content (52.3%) were desirable for high-level expression of the vaccine in bacteria. Considering the direct interaction between RBD protein and the ACE2 expressing cells, RBD immunization could induce specific neutralizing antibodies that may block this binding. Therefore, it can effectively hinder the invasion of the virus, making it more appropriate for vaccine development. For instance, researchers have been demonstrated that recombinant RBD contains multiple neutralizing epitopes that can induce a high titer of neutralizing antibodies (nAbs) against SARS-CoV (40). Eventually for future research, it is suggested further validation of our immuno-informatics results, the peptide expression in a bacterial host (E.g. BL21) and fulfill the several immunological assays is essential for more research in the future as a promising SARS-CoV-2 vaccine candidate.

